# Flexible open-source automation for robotic bioengineering

**DOI:** 10.1101/2020.04.14.041368

**Authors:** Emma J Chory, Dana W Gretton, Erika A DeBenedictis, Kevin M Esvelt

## Abstract

Liquid handling robots have become a biotechnology staple^1,2^, allowing laborious or repetitive protocols to be executed in high-throughput. However, software narrowly designed to automate traditional hand-pipetting protocols often struggles to harness the full capabilities of robotic manipulation. Here we present Pyhamilton, an open-source Python package that eliminates these constraints, enabling experiments that could never be done by hand. We used Pyhamilton to double the speed of automated bacterial assays over current software and execute complex pipetting patterns to simulate population dynamics. Next, we incorporated feedback-control to maintain hundreds of remotely monitored bacterial cultures in log-phase growth without user intervention. Finally, we applied these capabilities to comprehensively optimize bioreactor protein production by maintaining and monitoring fluorescent protein expression of nearly 500 different continuous cultures to explore the carbon, nitrogen, and phosphorus fitness landscape. Our results demonstrate Pyhamilton’s empowerment of existing hardware to new applications ranging from biomanufacturing to fundamental biology.

## MAIN TEXT

Automation has been widely implemented in biotechnology^3^ to facilitate routine tasks involved in DNA sequencing^4^, chemical synthesis^5^, drug discovery^6^, and molecular biology^7^. In principle, flexibly programmable robots could enable diverse experiments beyond the capabilities of human researchers, across a range of disciples within the sciences. Existing robotic software easily automates protocols designed for hand pipettes, but struggles to enable more specialized or sophisticated methods. As such, truly custom robot manipulation remains out of reach for most laboratories^2^, even those with well-established automation infrastructures.

Bioautomation lags behind the rapidly advancing field of manufacturing, where robots are expected to be task-flexible, responsive to new situations, and interactive with humans or remote management systems when ambiguous situations or errors arise^2^. A key limitation is the lack of a comprehensive, suitably abstract, and accessible software ecosystem^8–10^. Though bioinformatics is becoming increasingly open-sourced^11,12^, bioautomation has been slow to adopt key practices such as modularity, version control, and asynchronous programming.

To address these issues, we developed Pyhamilton, a Python package that not only facilitates high-throughput operations within the laboratory, but also allows liquid-handling robots to execute previously unimaginable and increasingly impressive methods. With this package, users can use process scheduling, run simulations for experimental planning, implement error handling for straightforward troubleshooting, and easily integrate robots with external laboratory equipment.

### Design of Pyhamilton Software

Pyhamilton enables Hamilton STAR and STARlet liquid handling robots to be programmed using standard Python. This allows for robotic method development to benefit from standard software paradigms, including exception handling, version control, object-oriented programming, and other cornerstone computer science principles (Supplementary Table 1). Pyhamilton seamlessly connects with Hamilton robots, can interface with custom peripherals (Fig. 1A), and contains unique Python classes corresponding to robotic actions (i.e. aspirate and dispense) and consumables (i.e. plates and pipette tips). To enable method troubleshooting, Pyhamilton can also simulate methods through Hamilton run control software and incorporate any Python package (i.e. enabling error notifications via push, text message, or Slack). Finally, in addition to the functionalities we present, researchers can now also develop their own flexible code that may be useful for increasingly specialized applications.

**Figure 1:**
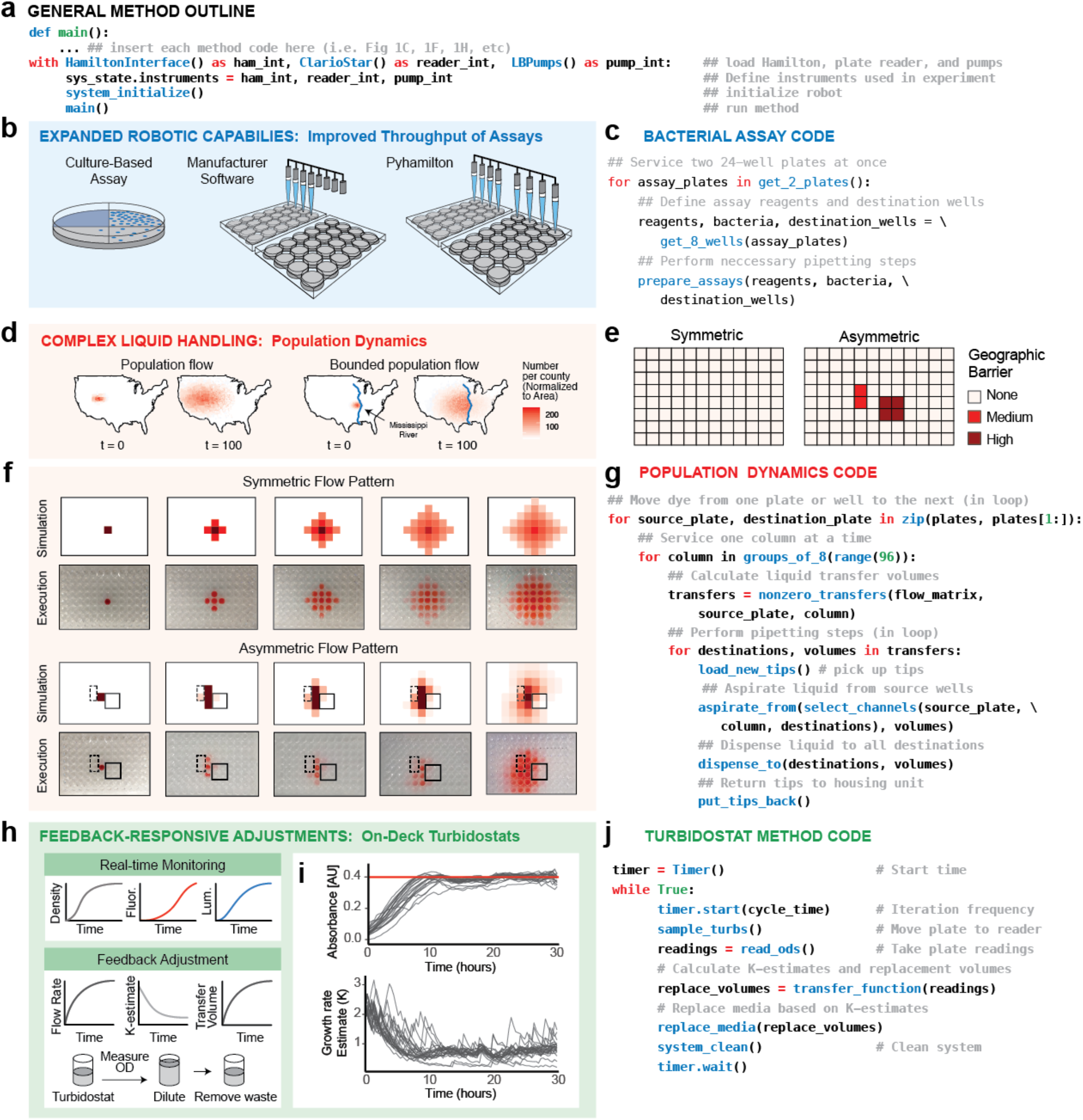
Example Pyhamilton Applications. **(a)** Generalizable Python outline for writing custom Pyhamilton code to interface with robot and integrated equipment such as plate readers (e.g., ClarioStar) and custom pump arrays. **(b)** Expanded robot capabilities allow for improved throughput of laboratory across 24-well plates. **(c)** Example code required to run a bacterial assay across multiple simultaneous plates. Code for bacteriophage plaque assay show (see supplemental methods). **(d)** Implementing complex and arbitrary bi-directional liquid handling to simulate experiments such as unbounded (left) or bounded (right) population flow across a geographic region, such as a river. **(e)** Geographic “barriers” described in matrix format **(f)** Simulation of bounded and unbounded migration (top), and visualization of the liquid patterns executed by the robot each iteration (bottom). Solid box designates “high” geographic barrier, dashed box designates a “medium” geographic barrier. **(g)** Example code required to run population dynamics simulations, using a sparse matrix to assign source wells, destination wells, and volume transfer fractions. **(h)** Real-time monitoring of on-deck turbidostats enables feedback control to equilibrate cultures to a set density. **(i)** Plate reader measurements for OD (top), and respective estimated growth estimates (bottom) obtained from data from 24 replicates. Data are smoothed with rolling mean and outlier points are excluded. OD set-point shown in red. **(j)** Example code required to maintain on-deck turbidostats using a transfer function to calculate k-estimates and volume transfer rates.

### Enabling improved throughput of laboratory assays

Standard liquid-handling software limits access to the full physical capabilities of a pipetting robot. For example, an 8-channel head cannot be readily programmed to pipette into two 24-well plates simultaneously although doing so is physically possible (Fig. 1B). This limits the ability to automate many laboratory assays in higher throughput: automation of methods involving 24-well plates is no faster than hand-pipetting, since both a robot and researcher can only pipette a single plate at a time. Thus, we first used Pyhamilton to develop a method which pipettes liquids over two 24-well plates simultaneously (Fig. 1C), doubling the speed. This can be critical for bacterial assays involving heated liquid agar which solidifies quickly. This simple example demonstrates the advantages of making full use of the robot’s mechanical capabilities, freed from software constraints.

### Enabling liquid transfers requiring complex calculations

Despite having far greater physical capabilities than a fixed-volume multi-channel pipette, it is difficult to implement complex liquid transfer patterns on a robot because programming using standard software is prohibitively monotonous. The ability to faithfully execute experiments involving hundreds of different pipetting volumes could enable new types of applications such as evolutionary dynamics experiments examining gene flow^13^, population symbiosis^14^, sources and sinks^15^, genetic drift^16,17^, and the spread of gene drive systems^18,19^ (Fig. 1D). We accordingly used Pyhamilton to enable the flexible transfer of organisms between populations in a 96-well plate, using preprogrammed migration rates to simulate geographic barriers (Fig. 1E).

A human would have great difficulty performing or programming hundreds of variable pipetting actions in many directions, in any reasonable time frame, without errors. With Pyhamilton, simple abstractions and data structures make this task straightforward. Instead of exhaustively specifying each pipetting step, we specified liquid transfer patterns as matrices, and allowed the software to compile the requisite steps. We demonstrate liquid transfer to nearby plates and between adjacent wells to model “flow” or “diffusion” across the miniaturized landscape of a 96-well plate. We then simulate genetic flow by visualizing the point spread of a drop of dye near the center of a plate (Fig. 1F). The amount of liquid exchanged and the number of wells is arbitrary, defined as a sparse matrix where the rows are source wells, the columns are destination wells, and the values are the fraction of liquid transferred (Supplementary Figure 2). Each iteration, the robot performs several hundred bi-directional liquid transfers to apply the matrix operations. Succinct code (Fig. 1G) can generate both symmetric and asymmetric diffusion patterns, which could be combined with a phenotypic reporter to experimentally simulate arbitrarily directionally bounded or unbounded migration (Fig. 1D) with many model organisms such as *E. coli,* yeast, or even nematodes.

### Enabling feedback control to maintain turbidostats

Though most liquid handling robots are used to execute a list of precompiled instructions (e.g., assembling reagents for many PCRs), many potential applications require making real-time modifications. For example, a turbidostat is a culture of cells that is maintained at a constant density by making real-time adjustments to the flow rate of media in response to turbidity sensing. In practice, this is accomplished with process controls which measure the optical density (OD) of a culture *in situ.* However, turbidity probes are both costly and not amenable to high throughput^20,21^. Thus, we sought to leverage the flexibility of Pyhamilton to multiplex the maintenance of many bacterial turbidostats by adjusting the volume of liquid transfers in response to real-time density measurements obtained using an integrated plate-reader (Fig. 1H). The method equilibrates each culture, growing in a multi-well microplate, to a set point (Fig. 1I) in response to these measurements by applying a transfer function to calculate the growth rate (k-value) and adjustment volume for each individual well over time (Fig. 1J).

### Asynchrony enables high-throughput turbidostats

To maximize the number of turbidostats that can be maintained, we next developed a more complex method which uses asynchronous programming to execute multiple robotic steps simultaneously— in this case plate reading and pipetting (Supp Fig. 1). This allows for nearly 500 cultures to be maintained with real-time fluorescent reporter monitoring on a single robot. In this method, bacterial cultures are inoculated into 96-well clear-bottom plates and their ODs and fluorescence levels are measured with an integrated plate reader (Fig. 2A). To minimize waste, consumables, and prevent media contamination, we also implemented a cleaning process (Fig. 2A): after each media transfer, each tip is sterilized with 1 % bleach, rinsed in water, and returned to its housing unit (Fig. 2A). To further minimize the possibility of cross-contamination between wells, each culture is assigned its own tip and media reservoir by housing replenishing media within high-volume 96-well plates. We confirmed that this method introduces no measurable cross contamination by inoculating 96 turbidostats with four different bacterial cultures expressing RFP, YFP, CFP, or no fluorescent protein in a gridlike pattern with no-bacteria controls (Fig. 2C). We then monitored the absorbance and fluorescence levels in real-time, and maintained the cultures at OD 0.8 for 24 hours. We observed no cross-contamination and no growth in the no-bacteria controls (Fig. 2C). We also inoculated the same bacterial strains at 6 different starting densities (OD=0.0-0.8) and demonstrated that irrespective of initial conditions, the feedback control algorithm equilibrates each culture to its set point within 12 hours (Fig. 2D).

**Figure 2:**
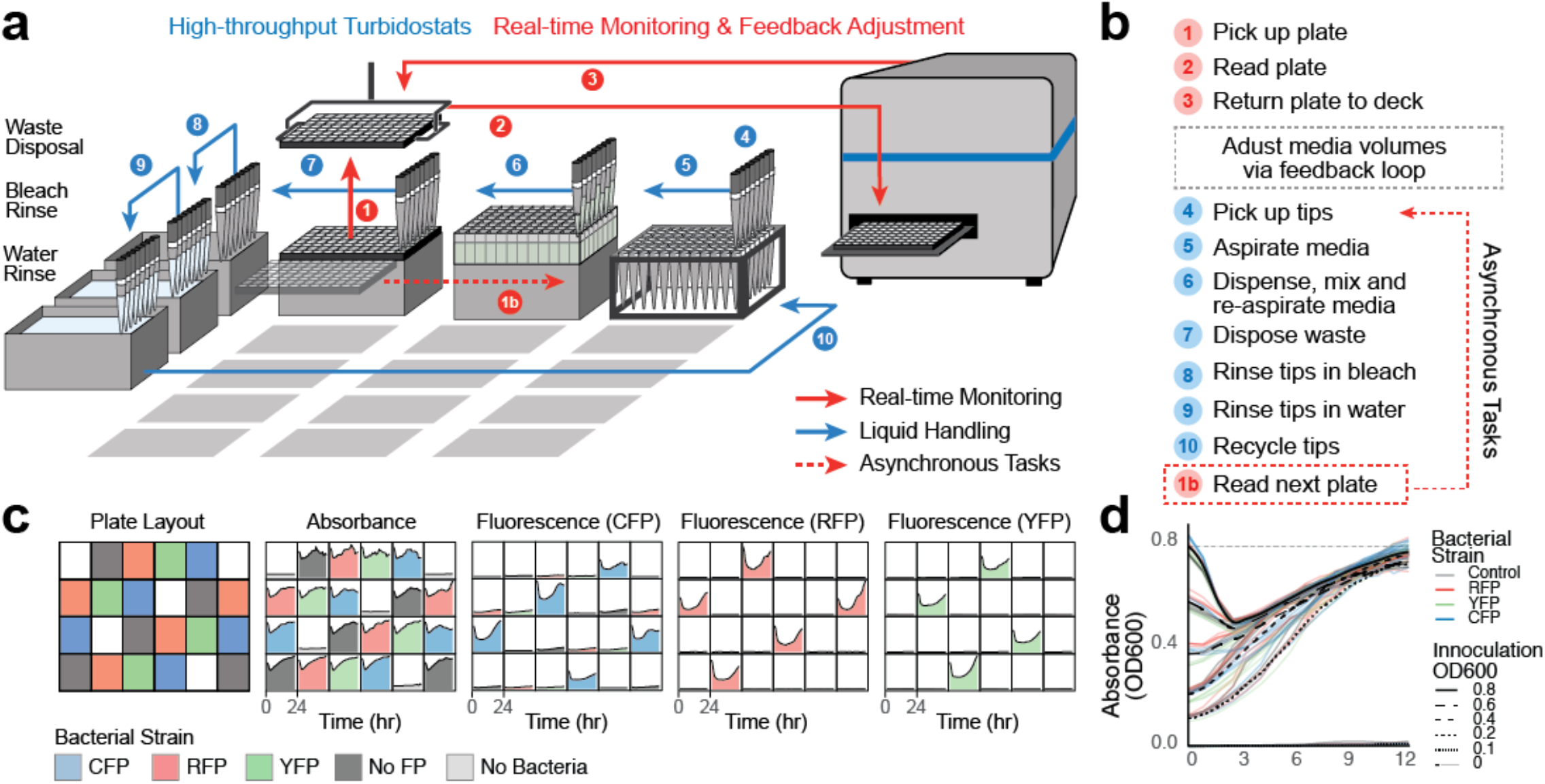
High-throughput turbidostats. **(a)** High-throughput turbidostat summary for up to 480 simultaneous evolutions. Bacterial populations are housed in 96-well clear-bottom plates on the deck of a liquid handling robot. Liquid handling is used to create a turbidostat in every well, continuously refreshing each population by diluting the bacterial culture from a respective deep-well media reservoir on deck. An integrated plate reader is used to monitor absorbance, luminescence, or fluorescence readouts for each culture. Movements by robotic pipette (blue arrow) and plate reader (red arrow) are shown. Dotted lines indicate tasks that are executed asynchronously, and require 10 minutes per plate. **(b)** Step-by-step summary of high-throughput turbidostat method. **(c)** Plate layout of real-time absorbance, CFP, RFP, and YFP fluorescence readings of 96 simultaneous cultures inoculated with either no bacteria, FP-null bacteria, and CFP, RFP, or YFP-expressing bacteria. Data shown from 24 representative wells. **(d)** Real-time absorbance measurements of 96 cultures inoculated at ODs of 0, 0.1,0.2 0.4, 0.6, 0.8, which equilibrate to a set point of 0.8 within 12 hours, consistent with simulation (Supplemental Figure 3).

### High-throughput perturbation analysis of metabolites

We next sought to use high-throughput turbidostat tracking to address an outstanding question in metabolic engineering by systematically mapping the chemical landscape that supports bacterial growth and protein expression. To do this, we surveyed the contributions of carbon, nitrogen, and phosphorus on growth and recombinant protein production by permuting chemical gradients for these metabolites in high-throughput. This effort, while seemingly well-studied, is difficult to accomplish without the proper number of replicates, experimental controls, long-term maintenance of log phase growth, and real-time monitoring, each of which are trivial to implement with Pyhamilton.

It has traditionally been thought that cells regulate protein production by allocating their resources to optimize for both expression and growth^22,23^. However, it has recently been shown that in either carbon-, nitrogen- or phosphorus-limiting conditions, cells are able to fine-tune their ribosomal usage to maintain equal levels of protein^24^. Thus, we hypothesized that exploration of the entire metabolite landscape (Fig. 3A) could more rigorously identify bacterial growth conditions optimized for recombinant protein production. To do this, we inoculated cultures with *E.coli* BL21, a strain commonly used for recombinant protein production in metabolic engineering or biomanufacturing, engineered for high constitutive expression of a fluorescent protein (CFP)^25^.

In a single experiment spanning 36 hours with no user intervention, we simultaneously quantified the equilibrium logphase growth rates and respective fluorescence levels of 300 individual turbidostats, representing 100 different media compositions in triplicate (Figure 3B). Cells were grown in modified M9 media containing 100 different ratios of carbon, nitrogen, and phosphorus and the cultures were maintained in log phase growth for 36 hours with feedback control (Supplemental methods). All cultures grew within +/− 20% of M9 media growth rate, with the exception of cultures that were starved of both carbon and phosphorus (Fig. 3C). We observed that increases in growth rate are primarily correlated with increases in phosphorus (independent of nitrogen or carbon levels), which is likely a result of increased DNA synthesis. Further, in phosphorus-limiting conditions, we find that the depressed growth rate can be rescued by supplementing carbon, but not nitrogen, suggesting that carbon precursors are a more limiting reagent than amino acids in metabolism (Fig. 3C).

**Figure 3:**
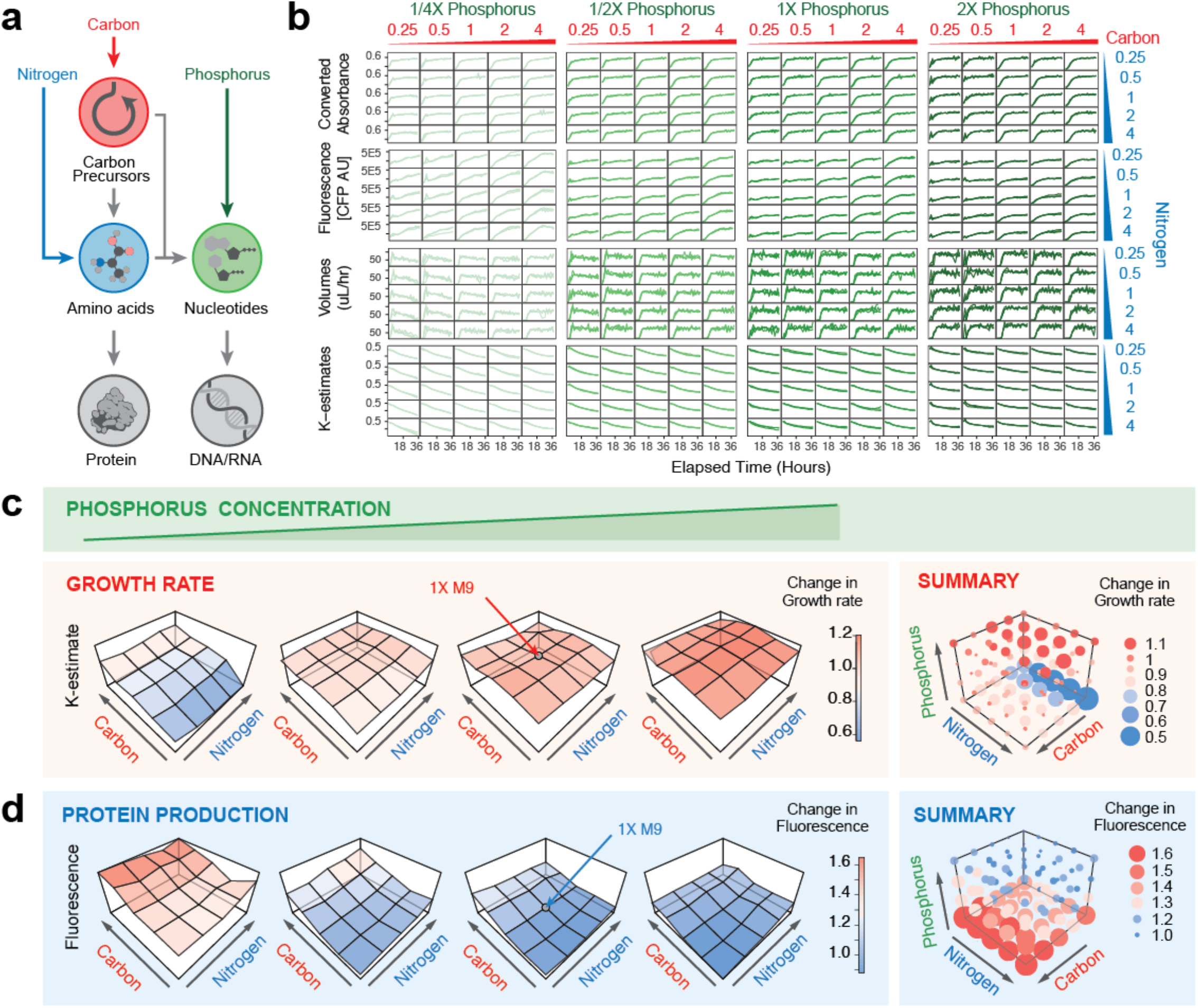
Metabolic profiling of protein production. **(a)** Schematic flow of carbon, nitrogen and phosphorus nutrients in protein and nucleotide production. **(b)** Real-time absorbance and fluorescent reporter monitoring for 100 various M9 media compositions (n=3 per condition). Real-time calculations of volumes/hr and estimates for k-value convergence shown. **c)** (left) Average growth rate for each media composition plotted as a 2dimensional fitness landscape of carbon and nitrogen, for four concentrations of phosphorus. (right) Summary of all 100 conditions shown as 3D fitness landscape colored by growth rate (blue = low, red = high). Size of dot indicates absolute deviation from average 1X M9 media composition. **d)** (left) Average amount of protein expression (measured by fluorescence) of each media composition plotted as a 2-dimensional fitness landscape of carbon and nitrogen, for four concentrations of phosphorus. (right) 3D protein-production landscape of all 100 conditions colored by amount of fluorescence (blue = low, red = high). Size of dot indicates absolute deviation from average 1X M9 media composition.

Consistent with previously published results^24^, we observe that the total amount of protein is generally not affected by limiting carbon or nitrogen, nor by supplementing the cells with excess of either nutrient. However, perhaps most interestingly, we additionally find that when phosphorus is limited (0.25X), excess carbon supplementation not only rescues the growth rate of the culture (Fig. 3C), but also results in an increase in total fluorescence (Fig. 3D). Since we observe minimal growth defects in these conditions, this finding suggests that on a per-cell basis, supplementing carbon in phosphorus-limiting conditions (such as in the soil^26,27^ or P-limited lakes^28^) can shunt bacterial metabolism from DNA/mRNA synthesis to protein translation without sacrificing growth. Collectively, these findings demonstrate that Pyhamilton enables researchers to answer rigorous metabolic engineering questions by enabling facile, low-consumable, yet rich hypothesis-generating experiments.

## DISCUSSION

Liquid handling robots have traditionally automated workflows that were explicitly designed for human researchers. Future methods enable experiments that could never be done by hand, such as protocols that must pipette continuously for multiple days, that perform complex calculations about future steps based on realtime data, or that make use of hardware that is more sophisticated than any hand-held multichannel pipette. Pyhamilton is an opensource Python framework which enables these types of experiments.

We showcase these improved capabilities by simultaneously quantifying the metabolic fitness landscape of 100 different bacterial growth conditions to identify ideal conditions for recombinant protein production. Though recent fluidic advances have enabled the maintenance of many continuous cultures^20^, the incorporation of real-time reporter monitoring vastly expands the types of questions that can be approached with facile, multiplex solutions. For example, one could maintain cultures of, and accurately quantify any reporter output for massively-parallel experiments including genetic knockout or CRISPR collections^29,30^, mutagenesis variants^31^, or even small-molecule compound libraries^32^. With high accuracy, any suspension culture of mixed populations could be maintained in log phase growth for days in order to study transient invaders into microbial communities^33^ or even microbiome system dynamics^34^. The advent of small molecule fluorescent reporters for metabolic fitness^35^, pH^36,37^, and CO_2_^38^, in addition to the hundreds of fluorescent protein sensors available to the synthetic biology community at large^39,40^, also impresses the seemingly unlimited potential of being able to multiplex and quantify changes in growth, gene expression, and the environment in real-time.

As such, Pyhamilton is a small part of an ongoing transition to a paradigm which leverages insights from computer science^8^ and applies them to biology. Similar to how Bioconductor^11^ and The Biopython project^12^ have revolutionized computational biology, bioinformatics, and genomics, our hope is that by making this software open-source and freely available, a community of scientists and developers could begin to similarly transform bioautomation. The experiments we have described represent only a small sampling of many possible Pyhamilton applications. Collectively, they highlight the potential of high-throughput robotic systems to transcend the repetitive processes for which they were conceived and directly address broad questions in microbiology, genetics, and evolution that are beyond the physical capabilities of human researchers.

## ACKNOWLEDGEMENTS

We would like to acknowledge Alvaro Cuevas of Hamilton Robotics for his examples, guidance, and assistance in making use of the Original Equipment Manufacturer (OEM) interface, along with the rest of Hamilton Robotics. We thank Jason Yang, Stephen Von Stetina, Ethan Alley, Brian Wang, Samantha Shepherd, and Timothy Erps for thoughtful comments and discussion.

## AUTHOR CONTRIBUTIONS

*Software:* DWG*. *Conceptualization:* EJC, DWG, EAD, and KME. *Methodology:* EJC, DWG, EAD. *Validation:* EJC, DWG, EAD. *Formal Analysis:* EJC. Investigation: EJC, DWG. Writing – Original Draft: EJC. Writing – Review & Editing: EJC, DWG, EAD, and KME. *Visualization:* EJC. *Funding Acquisition:* KME.

*All correspondence regarding Pyhamilton software development should be directed to DWG: (, https://github.com/dgretton/).

## METHODS

### Robotic Equipment set-up and interfacing

A Hamilton Microlab STARlet 8-channel base model was augmented with a Hamilton CO-RE 96 Probe Head and a Hamilton iSWAP Robotic Transport Arm. Air filtration was provided by an overhead HEPA filter fan module integrated into the robot enclosure. A BMG CLARIOstar luminescence multi-mode microplate reader was positioned inside the enclosure, within reach of the transport arm. *Software.* A general-purpose driver method was created using MicroLab STAR VENUS ONE software and compiled to Hamilton Scripting Language (hsl) format. Instantiation of this method and management of its local network connection was handled in Python. The Pyhamilton Python package provided an overlying control layer interface to the CLARIOstar plate reader in supporting Python packages. We used Git to develop and version control the packages and the specific Python methods used for each experiment; our software implementation can be found on github at https://github.com/dgretton/pyhamilton.

### Bacterial assays

For bacterial assay validation, bacterial plaque assays were used to confirm dilutions and agar solidification. Briefly, overnight cultures of S2060 cells were grown in 2XYT media supplemented with maintenance antibiotics were diluted 1,000-fold into fresh 2XYT media with maintenance antibiotics and grown at 37 °C with shaking at 230 rpm to OD_600_ ~0.6–0.8 before use. M13 bacteriophage were serially diluted 100-fold (4 dilutions total) in H_2_O. 20 μL of bacterial were added to 100 μL of each phage dilution, and to this 200 μL of liquid (70 °C) “soft” agar (2XYT media + 0.6% agar) supplemented with 2% Bluo-Gal was added onto a well of a 24-well plate already containing 235 μL of hard agar per well (2XYT media + 1.5% agar, no antibiotics). To prevent premature cooling of soft agar, the soft agar was placed on the robot deck in a 70 °C heat block. After solidification of the top agar, plates were incubated at 37 °C for 16–18 h. Source code from our implementation can be found at: https://github.com/dgretton/roboplaque

### Population Dynamics Experiments

Briefly, 96-well clear-bottom plates were filled with 100 uL of water in each well. Nucleation was initiated by adding colored dye to the first well, and liquid transfers were initiated and compiled by a Hamilton Microlab STARlet. Source code from our implementation can be found at: https://github.com/dgretton/pyhamilton_population_dynamics

### Feedback controller algorithm

Bacteria optical density (OD) was modeled to evolve as:

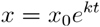

where *x* is the culture OD, *x*_0_ is the initial OD, *k* is the bacteria exponential growth constant (k-value) in reciprocal hours, and *t* is elapsed time in hours. A media replacement cycle is modeled as dilution of a culture by instant uniform mixing with transparent media of a fraction *y* of its initial volume, which linearly scales its OD *x* to a new OD *x*’ (e.g. if a 100 μL culture is at OD 0.3 and 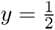, then the replacement is modeled as diluting with 50 μL transparent media, and the final OD *x*’ is 0.2), summarized as:

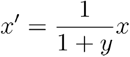

The culture OD is to be maintained at a constant setpoint, *x^set^*. In each cycle ^i^ = (),…, each representing a time interval Δ*t*, the turbidostat controller is responsible for producing an output command and state update according to a transfer function:

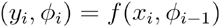

where *y_i_* is the new controller output command as a fraction of the turbidostat volume, *ϕ_i_* is the new controller internal state, *x_i_* is the present OD measurement, and *f*(*x_i_*, *ϕ*_*i*–1_) is the controller transfer function based on the OD measurement and the previous controller state *ϕ*^*i*–1^. The controller state may depend on the history of prior OD measurements *x*_0_,…, *x*_*i*–1_ and prior controller commands *y*_0_,…,*y*_*i*–1_.

#### Specific controller state

A feedback controller with a distinct state was created for each culture. The controller state is a triple 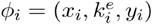 the present OD measurement, *x_i_*; the current estimate of the culture’s growth k-value, 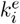; and the output command, *y_i_*.

#### Transfer function

The transfer function updates the three state variables and computes an output. Rearranging the model equations, we calculate the current k-value, given a new measurement *x_i_* taken an interval Δ*t* after the previous replacement executed, as

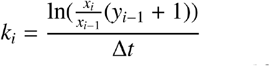

This *k* contributes to the state k-value estimate 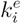 through a first-order linear filter to dampen the effect of measurement noise. The output to restore the turbidostat OD to the setpoint is

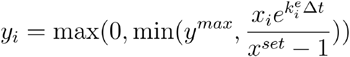

where the final output *y^i^* is subject to physical limits, being both nonnegative and not greater than the largest volume the robot can move with a pipette tip as a fraction of the turbidostat volume, appearing as *y^max^*. After output limiting, *y_i_* is saved in the controller state. Controller was developed as an abstract Python class and tested in simulation with mechanical and measurement noise models before application in experiments (Supplemental Figure 3). Filtered k-value estimates were used to draw conclusions about bacterial growth rates. Source code for implementation can be found at: https://github.com/dgretton/many_basic_turbidostats/blob/master/turb_control.py.

### On-deck Turbidostat cultures

#### Peristaltic pump array

To pump media onto the deck, up to seven miniature 12 volt, 60 mL/min peristaltic pumps (“fish tank pumps”) were actuated by custom motor drivers. A Raspberry Pi mini single-board computer received instructions over local IP and commanded the motor drivers via I^2^C (extended pump configuration details, see https://www.biorxiv.org/content/10.1101/2020.04.01.021022v1). Following each filling of the reservoir with fresh media, media was added to each bacterial turbidostat growing in a 24-well plate, based on OD and parameter estimation. Each turbidostat was then sampled by aspirating culture into a 96-well plate reader plate which was then read using an integrated ClarioStar plate reader. Remaining media was then drained from the reservoir and the system was rinsed 1X 5% bleach and 4X water between each iteration. S2060 bacterial strains were grown in 2XYT media supplemented with antibiotics. Source code for implementation can be found at: https://github.com/dgretton/many_basic_turbidostats.

### High-throughput Turbidostat cultures

#### Cell strains and growth conditions

To generate fluorescent reporter strains, plasmids pRSET-B YFP, pRSET-B mCherry, and pRSET-B mCherry were transformed into E. coli strain BL21(DE3) (New England Biolabs). Plasmids were a gift from Kalina Hristova (Addgene #108856, Addgene #108857, Addgene #108858). Bacteria cells were grown overnight in LB media, and the conditioned to grow in M9 Minimal Media: 33.7 mM Na2HPO4, 22.0mM KH3PO4, 8.5 mM NaCl, 9.35 mM NH4Cl, 0.4% Glucose, 1 mM MgSO4, 0.3 mM CaCl2, 1 ug biotin, 1 ug thiamin, 1X trace elements. For modified M9 Media, Phosphorus, Carbon, and Nitrogen sources were increased or decreased by 2 or 4 fold. For turbidostat inoculations, starter cultures were grown overnight at 37 °C for 16-18 h, and then diluted 1:100, and then grown for another 4-8 hours until in logphase growth. When each strain reached log-phase growth (OD 0.6-0.8), cultures were first diluted to an OD of 0.6 and then turbidostats were inoculated 1:100 into 175 uL in 96-well plate reader plates prior to initiation of the robotic method (Corning, Item#3631). Media for each well was aliquoted into a 96-deep well plate (Thomas Scientific, Item #1149J23). The robot deck was organized as described in Figure 2A. *Antibiotics.* Antibiotics (Gold Biotechnology) were used at the following working concentrations: carbenicillin, 50 μg/mL; chloramphenicol, 40 μg/mL. Source code for implementation can be found at: https://github.com/dgretton/many_asynchronous_turbidostats.

## Supplemental Figures

**Supplemental Figure 1:**
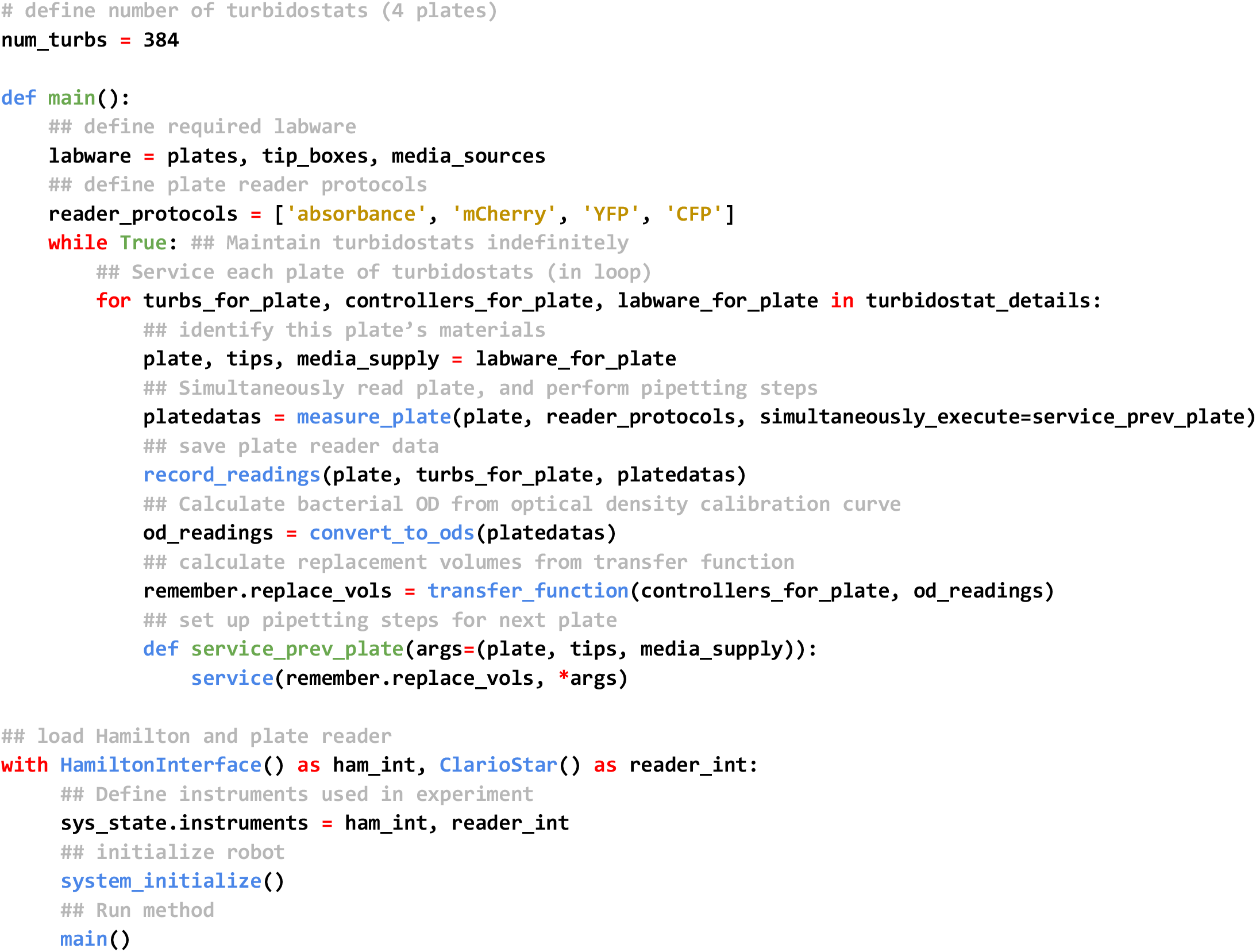
High-throughput turbidostats

**Supplemental Figure 2:**
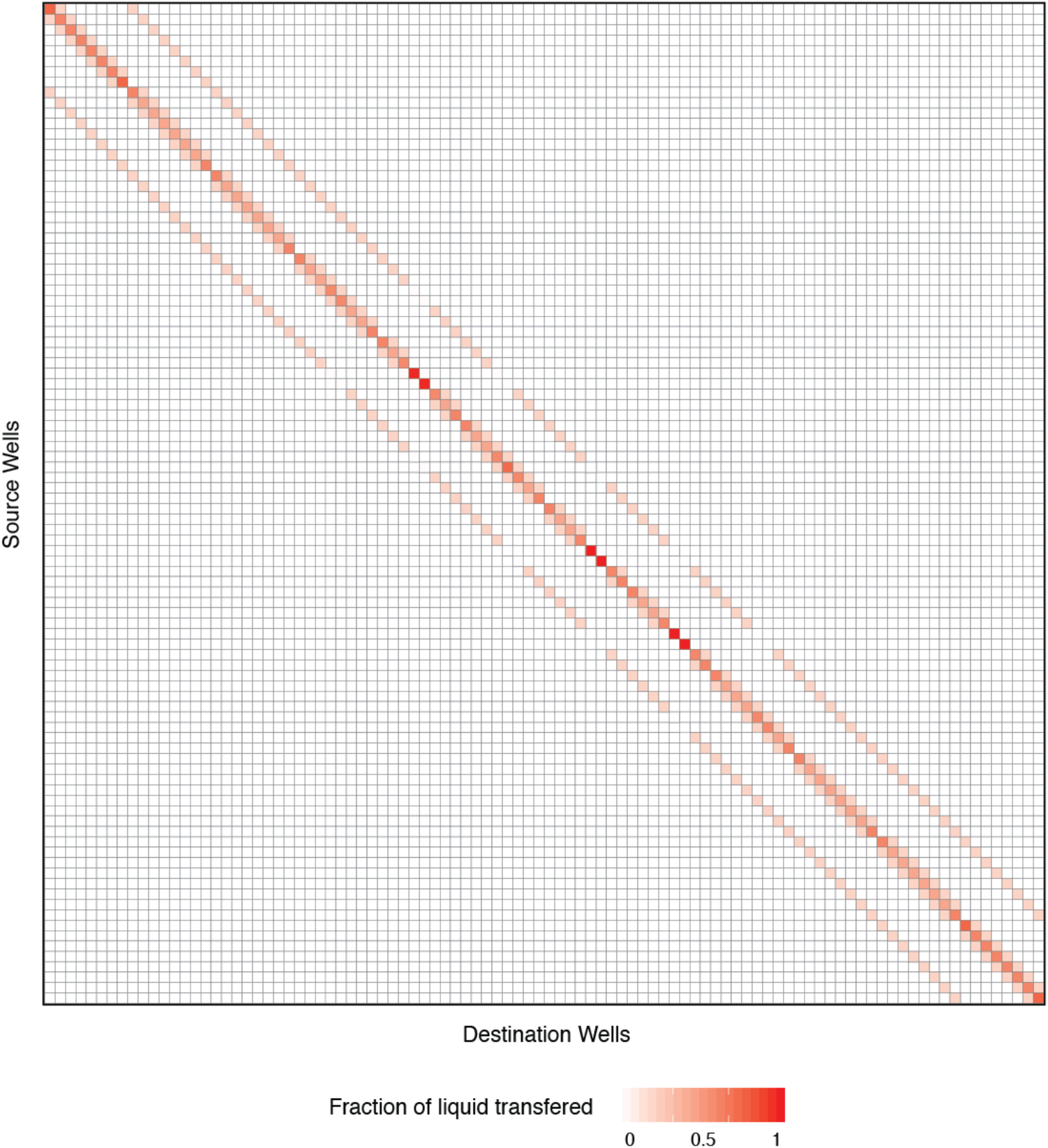
Population Dynamics Diffusion Matrix. Pyhamilton can be used in concert with all typical Python modules. The population dynamics application (Fig. 1D) makes use of matrix multiplication implemented by rectangular arrays from the scientific computing Python package NumPy. The diffusion matrix models arbitrary rates of flow from any of 96 wells in a microplate to all other wells. Gradient color scale represents fractional magnitudes of liquid transfers. Rows of the diffusion matrix sum to 1. Successive steps are computed by repeated multiplication of an initial 96-entry concentration vector by the 96×96 diffusion matrix. In this matrix, the dark main diagonal indicates that most of each population stays in its source well. Entries above and below the main diagonal by 1 and 8 rows represent transfers to vertical and horizontal neighbors respectively. All other entries are zero because population flow is not modeled to occur diagonally or beyond immediate neighbors. Obstacles to gene flow are captured by lighter colored areas in off-diagonal entries. Though this matrix is symmetric in that flows in both directions between each pair of wells are the same, matrix symmetry is not a requirement in general. The matrix construction facilitates offline analysis and visualization prior to robot execution. This construction would be difficult to implement in existing programming applications for Hamilton robots. Available online at: https://github.com/dgretton/pyhamilton_population_dynamics/blob/master/flow_matrix.csv.

**Supplemental Figure 3:**
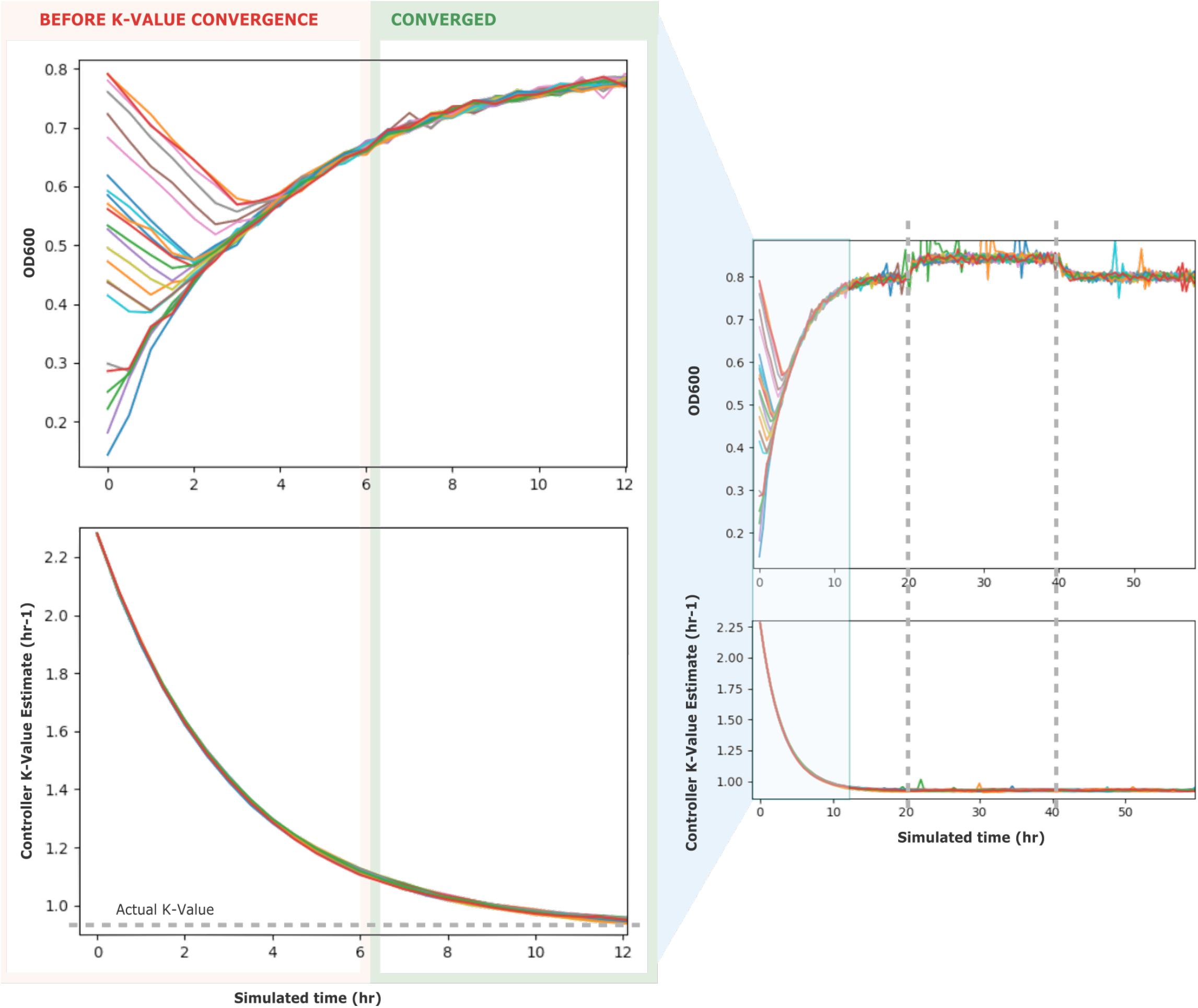
Turbidostat Controller Simulation. 24 simulated turbidostats using the same controllers as experiments with initial k-value estimate of 2.3 hr^-1^ (20-minute doubling time typical of E. Coli) and actual k-value 0.93 hr^-1^ (typical of metabolite-poor media) converge in 12 hours. Start conditions vary between OD 0.1 and OD 0.8. Simulation includes uniform pipetting volume noise model and power law measurement noise model (spurious peaks). Controllers initially over-replace media in higher density cultures due to growth rate overestimate, causing OD to drop temporarily, before recovering as the k-value estimate converges more closely to the actual k-value. This behavior is exactly recapitulated in the turbidostat convergence study (Fig. 2D), indicating good correspondence between model and system. The simulated turbidostat OD setpoint was adjusted ±0.1 at 20 hours and 40 hours. Though the OD measurements and the controller’s transfer volume commands both change at these times, the inferred k-value stays constant.

## Supplemental Tables

**Supplemental Table 1:** Pyhamilton Design Principles & Features

|  |  |
| --- | --- |
| Modularity | Use Pyhamilton alone as one complete package |
|  | Compatible auxiliary instrument modules can be used in concert or independently |
|  | Integratable as one part of a larger project |
|  | Expandable vocabulary of commands |
| Abstraction layers | Common interface for Hamilton Run Control simulator integration and physical robot |
|  | Independent of any particular robot model (STAR, STARlet, etc.) |
|  | Hierarchical exception definitions |
| Transparent connection management | Locally hosted background server |
|  | GUI-less launcher for background hsl executor |
|  | One-line context managers launch and destroy background processes automatically |
| Deployment readiness | Predefined semantic errors |
|  | Automatic command formatting with detailed defaults |
|  | Idiomatic support for parallel commands |
| Modern code practice | Legible code |
|  | Version control |

